# Loss of E3 ligase Ube4A Disrupts Colon Homeostasis and Accelerates Experimental Colitis via Altered Lipid Handling

**DOI:** 10.64898/2026.01.02.697430

**Authors:** Simon Guignard, Molee Chakraborty, Silvia Gonzalez-Nieves, David DeBruin, Emily Ebert, Anastasiia Vinogradskaia, Michelle Brennan, Ryan M Teague, Anutosh Chakraborty, Vincenza Cifarelli

## Abstract

**Background:** Ubiquitin-dependent signaling is essential for maintaining intestinal homeostasis and its dysregulation contributes to chronic intestinal disorders, such as Inflammatory Bowel Disease (IBD). Ube4A is a U-box E3/E4 ubiquitin ligase involved in lipid metabolism and insulin signaling in metabolic tissues. Autoantibodies against Ube4A have been identified in patients with IBD and are associated with disease long-term complications. Despite these clinical associations, the physiological role of Ube4A in the gastrointestinal tract remains unknown. This study aimed to define the function of Ube4A in the colon and determine how its loss influences susceptibility to experimental colitis.

**Methods:** *UBE4A* expression in human colonic tissue from healthy individuals and patients with IBD was analyzed using publicly available single-cell RNA sequencing datasets. The role of Ube4A in colonic homeostasis and colitis pathogenesis was examined using global Ube4A knockout (UKO) mice subjected to dextran sulfate sodium (DSS)-induced colitis. UKO colon phenotypes were characterized using transcriptomic analyses, immunofluorescence, and flow cytometry.

**Results:** *UBE4A* is highly expressed in human colonic epithelial cells, and its expression is reduced from healthy to IBD inflamed tissues. In mice, Ube4A deficiency significantly exacerbated DSS-induced colitis, as evidenced by increased weight loss, disease activity scores, shortened colon length, and more severe histological injury. Transcriptomic profiling revealed enhanced inflammatory signaling, alongside dysregulation of lipid transport and storage, as well as antimicrobial defense pathways. DSS-treated UKO mice also exhibited increased mast cell activation and elevated expression of matrix metalloproteinases. Importantly, colons from UKO mice displayed baseline transcriptional alterations indicative of epithelial stress and disrupted lipid metabolic programs, even in the absence of injury.

**Conclusions:** Ube4A is a previously unrecognized regulator of colon homeostasis. Its loss induces existing epithelial stress and metabolic reprogramming that sensitize the colon to exaggerated inflammatory responses during injury such as experimental colitis.

## INTRODUCTION

Ubiquitination is a reversible post-translational modification in which the small regulatory protein ubiquitin is covalently attached to substrate proteins through a highly conserved enzymatic cascade involving E1 activating, E2 conjugating, and E3 ligase enzymes [1]. Although ubiquitination classically targets proteins for proteasomal degradation, it also regulates protein stability, subcellular localization, and signaling, thereby controlling essential cellular processes including DNA repair, signal transduction, cell-cycle progression, and immune responses [2]. In the gut, ubiquitination is a critical regulator of barrier integrity and immune homeostasis, shaping epithelial responses to microbial and inflammatory cues [3]. Consistent with this role, increasing evidence shows that disruption of ubiquitin-mediated pathways can impair mucosal defenses and promote chronic intestinal inflammation [4].

Inflammatory Bowel Disease (IBD), which includes Crohn’s disease and ulcerative colitis, is a chronic, relapsing inflammatory disorder of the gastrointestinal tract with rising global prevalence. Although genetic, environmental, microbial, and immune factors contribute to disease susceptibility, the exact molecular mechanisms that initiate the disease remain elusive [5–7]. The lack of curative treatment for IBD contribute to progressive tissue injury and increase the risk of colorectal cancer and other severe complications [8, 9]. Accumulating data from both human studies and experimental preclinical models implicates dysregulated ubiquitin-dependent pathways in IBD pathogenesis [3, 4]. Genome-wide association studies have identified IBD risk variants in genes encoding ubiquitin-modifying enzymes [10–13], and several E3 ligases (e.g., RNF183, RNF20, A20, Pellino 3, TRIM62, and Itch) regulate key inflammatory pathways such as NF-κB signaling and ER stress responses during intestinal inflammation [14–22].

Clinical observations have highlighted Ube4A, a U-box E3 ubiquitin ligase, as a potential IBD-associated protein [23]. Autoantibodies against Ube4A have been detected in patients with Crohn’s disease, where their circulating levels were correlated with increased disease penetrance, complicated phenotypes (stricturing or penetrating disease), and elevated likelihood of surgical intervention [24]. Despite this clinical association, the biological function of Ube4A in the gastrointestinal system remains unexplored. Besides acting as a classic E3 ligase, Ube4A also exhibits a unique function, termed as E4 ligase activity that extends mono- or oligo-ubiquitin to poly-ubiquitin chains on protein substrates, influencing substrate fate and signaling dynamics [25]. Such regulatory capacity suggests that Ube4A may act as an integrative node in adaptive cellular programs. Yet, its cellular targets and tissue-specific roles remain largely unknown. A recently generated whole-body Ube4A knockout (UKO) mouse model revealed that Ube4A is an important regulator of metabolic health and lipid metabolism, particularly controlling lipid accumulation in liver and adipose tissue [26]. Studies show that perturbations in epithelial metabolic programs including mitochondrial energy metabolism and lipid handling, can directly influence epithelial regenerative capacity, barrier function, and susceptibility to chronic intestinal inflammation [27]. These finding raise the possibility that ubiquitin-dependent regulators of lipid metabolism such as Ube4A may exert important effects on intestinal homeostasis and inflammation. Here, we investigated the function of Ube4A in colonic tissue under steady-state conditions and in the context of Dextran Sodium Sulfate (DSS)-induced colitis, a widely used model that recapitulates key features of human IBD [28]. We used global UKO mice together with transcriptomic profiling, flow cytometry and confocal imaging to define how Ube4A loss remodels colonic epithelial homeostasis and susceptibility to injury and inflammation.

## RESULTS

### UBE4A Expression Patterns in Healthy and Inflamed IBD Human Colon

The physiological role of Ube4A in the intestine is largely unknown. To begin addressing this gap, we first examined its expression pattern in the human colon. Data from the Human Protein Atlas indicate that UBE4A is expressed in the colon of healthy individual [29]. To define its cellular distribution within the colon, we analyzed recently published single-cell RNA-sequencing (scRNA-seq) datasets from healthy and IBD-affected colon biopsies [30]. In healthy colon, *UBE4A* was predominantly expressed in epithelial cells and plasma B cells, with minimal expression in stromal compartments (**Figure 1A-C**). *UBE4A* expression in the epithelial compartment progressively declined from healthy to non-inflamed IBD to inflamed IBD colon (**Figure 1D**). Within the epithelial lineage, *UBE4A* mRNA was enriched in proliferating transit-amplifying (TA) progenitors (**Figure 1E**), which give rise to goblet cells, a lineage essential for mucin production and maintenance of the protective mucus barrier [30, 31].

**Figure 1.**
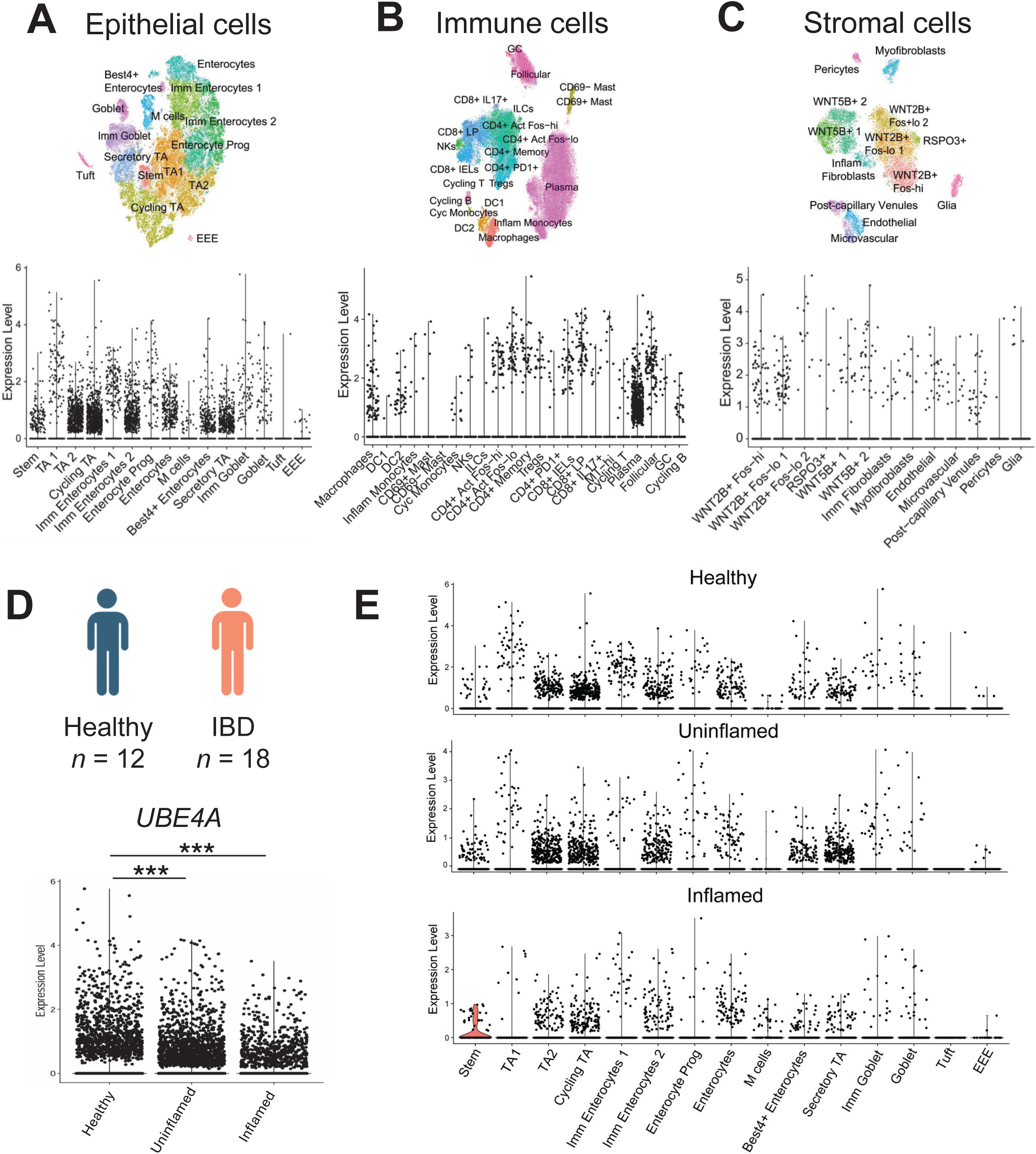
*UBE4A* is expressed in human colon and downregulated in IBD. Single-cell RNA sequencing analysis of *UBE4A* mRNA expression in healthy human colonic biopsies across major cellular compartments, including **(A)** epithelial, **(B)** immune, and **(C)** stromal cell populations. Dot plots showing *UBE4A* expression levels across annotated cell clusters within each compartment. **(D)** Comparison of *UBE4A* expression in scRNA-seq–derived epithelial cells from healthy controls (n=12) and IBD patients (total n=18), stratified into non-inflamed and inflamed colonic biopsies. **(E)** *UBE4A* expression across individual epithelial cell subpopulations in healthy, non-inflamed, and inflamed IBD colon. Each dot represents one individual cell. ***P < 0.001 by Wilcoxon rank sum test.

### Ube4A deletion exacerbates DSS-induced colitis in mice

Given the robust expression of UBE4A in the colon, its reduction in IBD, and the presence of Ube4A autoantibodies in patients with IBD, we next investigated whether loss of Ube4A contributes to impaired inflammatory response during colitis. To test this hypothesis, we employed oral dextran sulfate sodium (DSS) administration, a well-established chemical model of experimental colitis [28]. Sex- and age-matched wild-type (WT) and UKO mice were subjected to 2.5% DSS for five days to induce colon inflammation (injury phase), followed by 15 days of regular water (recovery phase) (**Figure 2A**). Compared to controls, UKO mice developed substantially more severe colitis, as indicated by a significant body weight loss (AUC: 102.9±2.4 for WT vs 79.9±5.3 for UKO) (**Figure 2B**) and a higher disease activity index (DAI) during the injury phase (AUC: 18.9±1.4 for UKO vs 10.5±0.6 for WT) (**Figure 2C**). After five days of DSS treatment, UKO mice also presented significant shorter colon length compared to WT (8.2±0.3 cm vs 6.8±0.4 cm), with evident presence of bleeding (**Figure 2D**). Histological examination of Hematoxylin and Eosin (H&E) staining revealed increased colonic damage in UKO colons (injury score: 9.9±0.8 vs 6.7±1.3), characterized by a higher prevalence of oedema (yellow arrow), immune cell infiltration (black arrow), and epithelial disruption or erosion (white arrow) (**Figure 2E-F**). To assess the impact of Ube4A deletion on the mucosal barrier, mucus-producing goblet cells were quantified using Alcian blue staining. Compared to WT mice, UKO mice showed a marked reduction in Alcian blue–positive goblet cells (**Figure 2G-H**). Together, these results indicate that loss of Ube4A worsens the severity of DSS-induced colitis.

**Figure 2.**
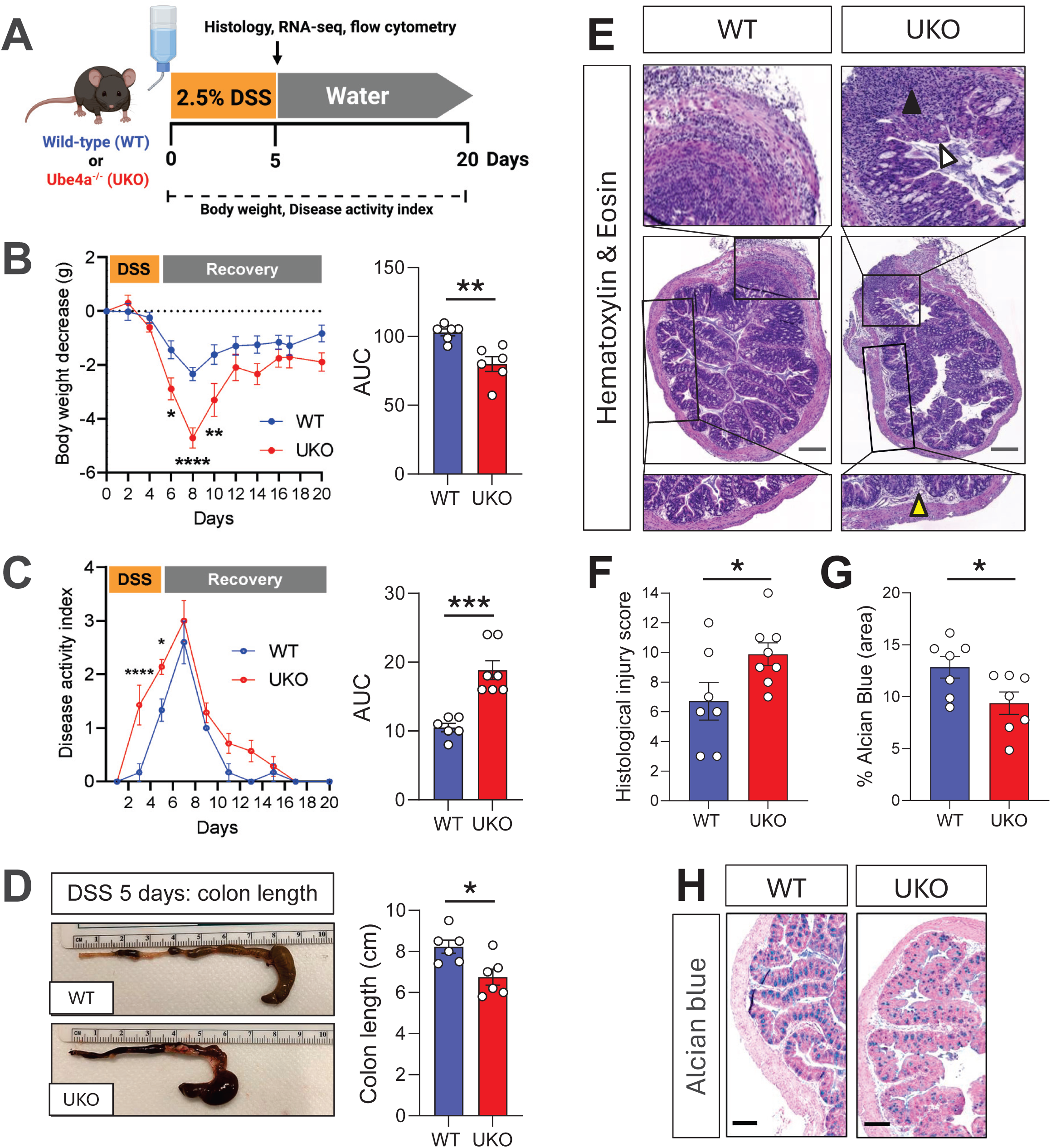
Ube4A deletion exacerbates DSS-induced colitis in mice. **(A)** Schematic of Dextran Sulfate Sodium (DSS)-induced colitis experimental model. Ube4A-deficient mice (UKO) or Wild-type (WT) mice were subjected to 2.5% DSS in drinking water for five consecutive days, followed by a recovery phase of 15 additional days in normal water. The severity of colitis in DSS-treated mice was assessed by measuring **(B)** body weight loss and **(C)** disease activity index (DAI) throughout the 20-day protocol. **(D)** Representative images and quantification of colon length (cm) after five days of DSS treatment. **(E)** Representative images of Hematoxylin and Eosin (H&E) staining of colon cross-sections. Scale bars: 500 μm. **(F)** Histological scoring of H&E staining based on three parameters: presence of oedema (yellow arrow), immune cell infiltration (black arrow), and epithelial disruption or erosion (white arrow), as described in Materials and Methods. **(G-H)** Alcian blue staining and quantification from five day-DSS-treated UKO and WT mice. Scale bars: 200 μm. Data are presented as mean ± SEM. Each dot represents one individual mouse (n=6-8/group). *P < 0.05, **P < 0.01, ***P<0.005 **** P < 0.0001 by two-way ANOVA, Mann-Whitney, and Student unpaired t-test.

### Ube4A deletion upregulates inflammatory and lipid storage pathways, while downregulating morphogenic pathways in DSS-treated mice

To identify the mechanisms contributing to the worsened colitis phenotype in UKO mice, we characterized their colonic transcriptomic signature by bulk RNA-sequencing after five days of DSS treatment. Compared to WT mice, we identified 663 differentially expressed genes (DEGs) in the colon of DSS-treated UKO mice. Among these DEGs, 457 were upregulated, while 206 were downregulated (**Figure 3A**). Functional enrichment analysis of the DEGs identified significant upregulation of several biological pathways related to inflammatory processes, lipid transport and microbial sensing, while biological pathways related to morphogenesis were downregulated (**Figure 3B**). Consistent with the pathway analysis, DSS-treated UKO colons exhibited a significant upregulation of DEGs involved in acute inflammatory response (*Cxcl9, Cxcl1, Ccl2, Il1b*), Th2 immunity (*Il1rl1, Gata3, Il5ra*), and lipid transport/storage (*Fabp1, Fabp2, Cidec, Mogat2*) (**Figure 3C-E**). In addition, several factors related to defense response against bacterium, including *Defb37*, *Reg3b*, and *Reg3g*, were also upregulated (**Supplementary Figure 1A**). Conversely, genes involved in morphogenesis (*Tnnt2, Tenm4, Actc1*) were downregulated in the colon of DSS-treated UKO mice (**Figure 3F**).

**Figure 3.**
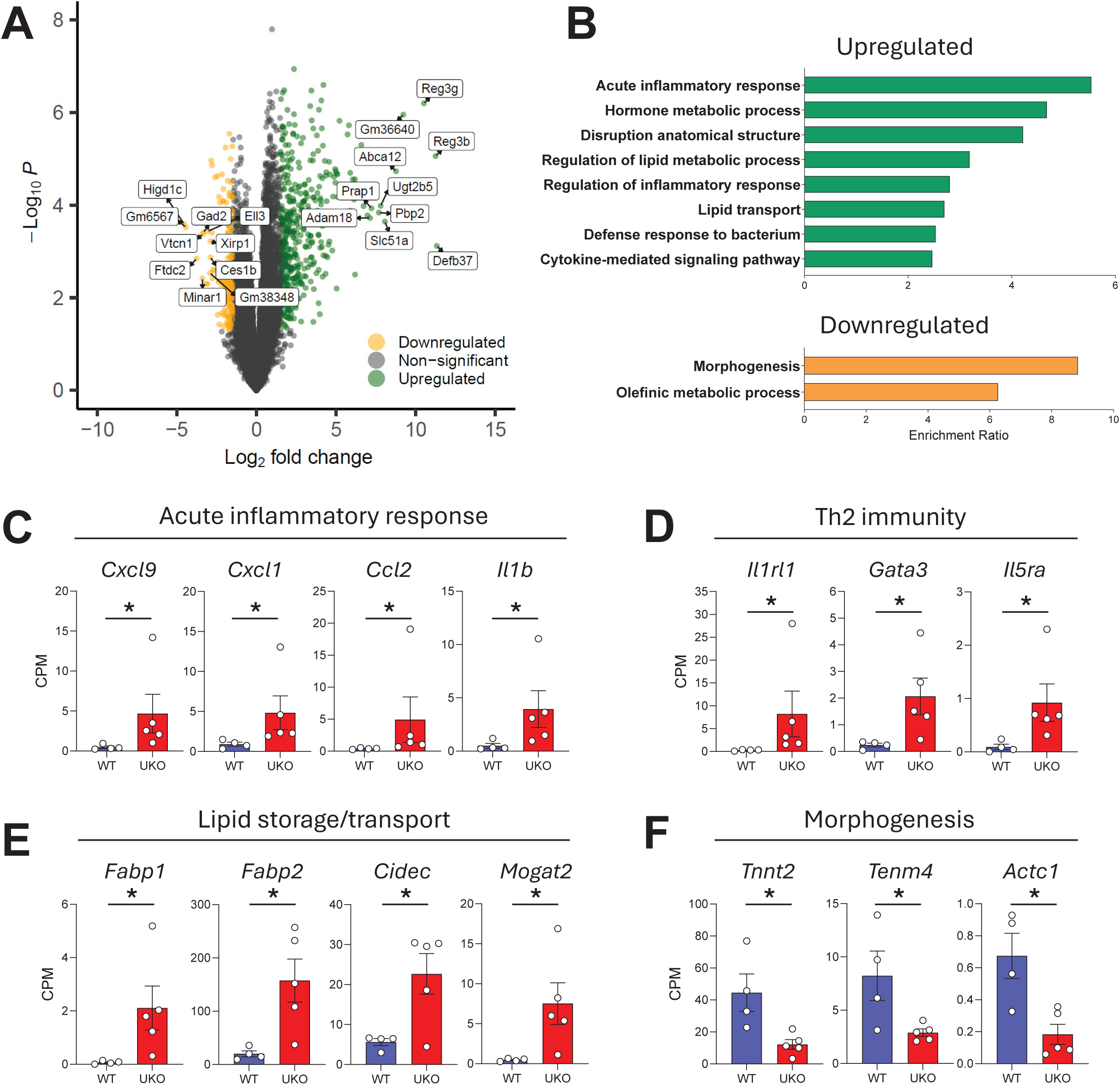
Loss of Ube4A promotes inflammatory and lipid storage transcriptional programs in DSS-induced colitis. Bulk RNA sequencing analysis of colon from five days DSS-treated WT and UKO mice. **(A)** Volcano plot showing significantly differentially expressed genes in DSS-treated UKO vs WT mice (Cut-off: p-value < 0.05 and log(fold change) < −1.5 or > 1.5), with upregulated genes shown in green and downregulated genes in orange. **(B)** Biological pathways enrichment analysis of upregulated (green) and downregulated (orange) DEGs. Relative expression of selected upregulated genes involved in **(C)** acute inflammatory response, **(D)** Th2 immunity, and **(E)** lipid storage/transport. **(F)** Downregulated genes related to morphogenesis. Data are presented as mean ± SEM. Each dot represents one individual mouse (n=4-5/group). *P < 0.05 by Benjamini-Hochberg.

### Ube4A deletion promotes colonic mast cell accumulation during experimental colitis

DSS-treated UKO colons showed a marked increase in the expression of Interleukin-1 receptor-like 1 (*Il1rl1*) (**Figure 3C**), which encodes the interleukin-33 receptor ST2— highly expressed on mast cells [32]. Consistent with this, immunofluorescence analysis revealed an increased expression of mast cell protease-1 (MCPT-1) staining, in the colons of DSS-treated UKO mice compared to WT controls (**Figure 4A–B**). Flow cytometry analysis further supported these findings, showing a higher frequency of mast cells (ST2⁺/c-kit⁺) in the mesenteric lymph nodes of DSS-treated UKO mice (**Figure 4C–D**). Since mast cell activation is closely linked to epithelial injury and extracellular matrix remodeling [33, 34], we next examined the expression of matrix metalloproteinases (MMPs), which play key roles in tissue degradation and repair during IBD [35]. Our RNA-seq analysis revealed that several MMPs were markedly upregulated in the colonic segments of DSS-treated UKO mice (**Supplementary Figure 1B**). Consistent with these findings, real-time PCR analysis confirmed the overexpression of transcripts encoding MMP3, MMP10, and MMP13 in the colons of DSS-treated UKO mice (**Figure 4E**).

**Figure 4.**
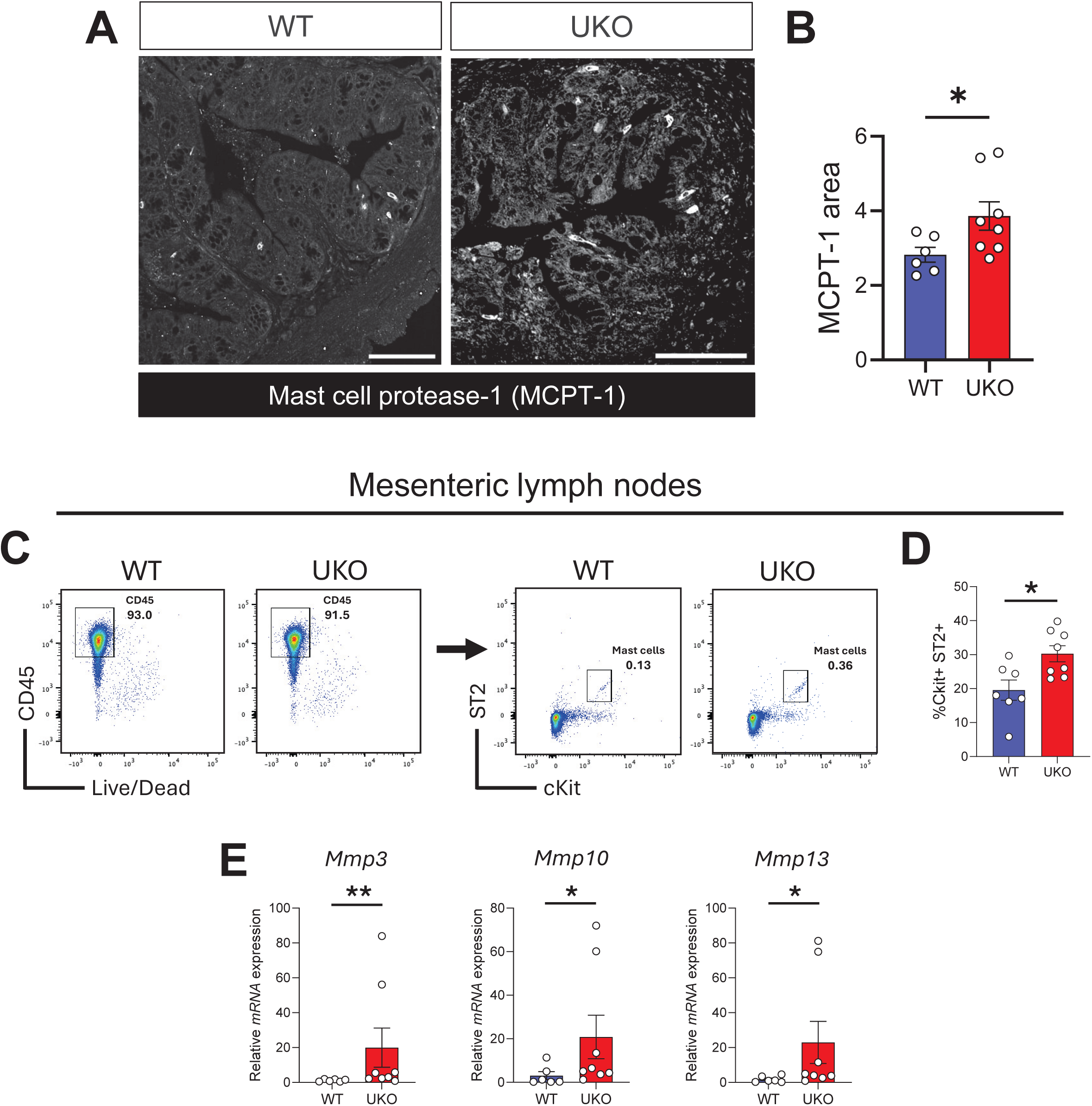
Ube4A deletion is associated with increased colonic mast cell activation and MMPs expression during experimental colitis. **(A-B)** Representative images and quantification of mast cell protease-1 (MCPT-1) staining of DSS-treated WT and UKO colons. Scale bars: 500 μm. **(C-D)** Flow cytometry analysis and quantification of mast cells (cKit+/ST2+) in DSS-treated WT and UKO mesenteric lymph nodes. **(E)** Relative expression of *Mmp3*, *Mmp10* and *Mmp13* in five-day DSS-treated WT and UKO colon by qRT-PCR. Data are presented as mean ± SEM. Each dot represents one individual mouse (n=6-8/group). *P < 0.05 by Mann-Whitney test.

### Metabolic dysregulation underlies epithelial stress in the Ube4A-deficient colon

To assess whether the increased DSS susceptibility in UKO mice arises from a pre-existing vulnerability, we next examined colonic homeostasis in untreated UKO mice. Consistent with previous reports, chow-fed UKO mice appeared healthy and maintained normal body weight [26]. H&E staining revealed no evident changes in the colon architecture and no gross signs of tissue damage (**Figure 5A**). Furthermore, Alcian blue staining showed no differences in goblet cell abundance or mucus production between WT and UKO mice (**Figure 5B-C**). To assess how loss of Ube4A influences colonic transcriptional programs, we next performed bulk RNAseq on colon tissues from unchallenged WT and UKO mice. As compared to WT, 570 genes were differentially expressed in colons from UKO mice (459 upregulated and 111 downregulated genes) (**Figure 5D**). Pathway analysis of the DEGs identified significant upregulation of biological pathways related to lipid transport (*Fabp1, Fabp2*, *Abca12, Prap1*) and isoprenoid metabolic process (*Aldh1a2, Hmgcs2*) (**Figure 5E-F**). In contrast, the downregulated pathways were related to complement activation (e.g., *C4bp*) (**Figure 5E, G**).

**Figure 5.**
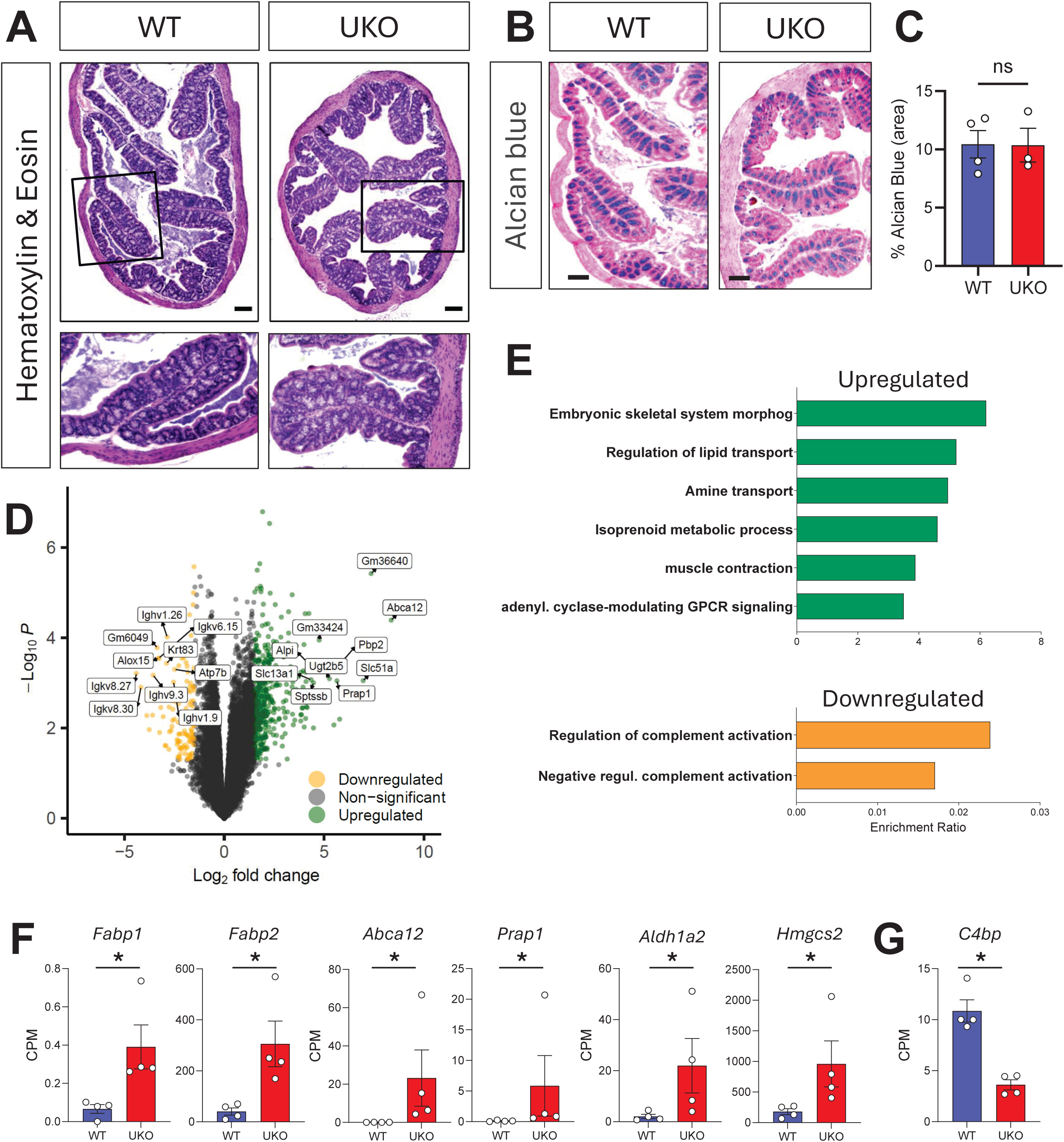
Ube4A deficiency triggers abnormal colonic transcriptional programs without altering histological tissue integrity in basal conditions. Representative images of **(A)** Hematoxylin and Eosin, and **(B-C)** Alcian blue staining in colons of unchallenged mice (this data included both male and female individuals). Scale bars: 200 µm. **(D)** Volcano plot showing significantly differentially expressed genes (DEGs) in unchallenged colons of UKO versus WT mice, analyzed by bulk RNA-seq (Cut-off: p-value < 0.05 and log(fold change) < −1.5 or > 1.5). **(E)** Biological pathways analysis showing top significantly upregulated (green) and downregulated (orange) pathways. Relative expression of selected **(F)** upregulated genes involved in lipid transport, isoprenoid metabolism, amine transport, and **(G)** downregulated genes related to complement activation. Data are presented as means ± SEM. Each dot represents one individual mouse (n=3-5/group). *P < 0.05, ns: not significant, by Benjamini-Hochberg or Mann-Whitney test.

To uncover molecular determinants that might predispose Ube4A-deficient mice to heightened susceptibility to colitis, we compared the colonic transcriptomes of UKO and WT mice under both unchallenged and DSS conditions. Strikingly, even in the absence of inflammatory challenge, UKO colons already displayed a marked altered transcriptional signature, with 194 genes consistently upregulated (**Figure 6A**) and only 32 downregulated genes across both conditions (**Figure 6D**). Gene ontology analysis showed that the shared upregulated genes were enriched in pathways related to lipid metabolic processes (including isoprenoid metabolism, steroid metabolic process, regulation of lipid metabolic process), with increased expression of key lipid-handling genes such as *Fabp1, Fabp2, Cidec,* or *Mogat2* (**Figure 6B-C)**. In addition, several epithelial stress- and damage-responsive genes, including *Reg3b, Reg3g, Hmgcs2*, *Plet1*, *Retnlb,* and *Vnn1* [36–39], were already elevated in UKO colons at baseline, as compared to controls (**Figure 6C** and **Supplementary Figure 2**). In contrast, among the most downregulated genes between baseline and DSS-treated colons (**Figure 6F**) were *Vtcn1*, an epithelial-derived inhibitory immune checkpoint [40] and *Atp7b*, a copper transporter linked to intestinal lipid transport and barrier function [41] (**Figure 6C** and **Supplementary Figure 2**). Together, these alterations suggest a potential immuno-metabolic role for Ube4A in the colon and may predispose Ube4A-deficient mice to intestinal injury upon challenge.

**Figure 6.**
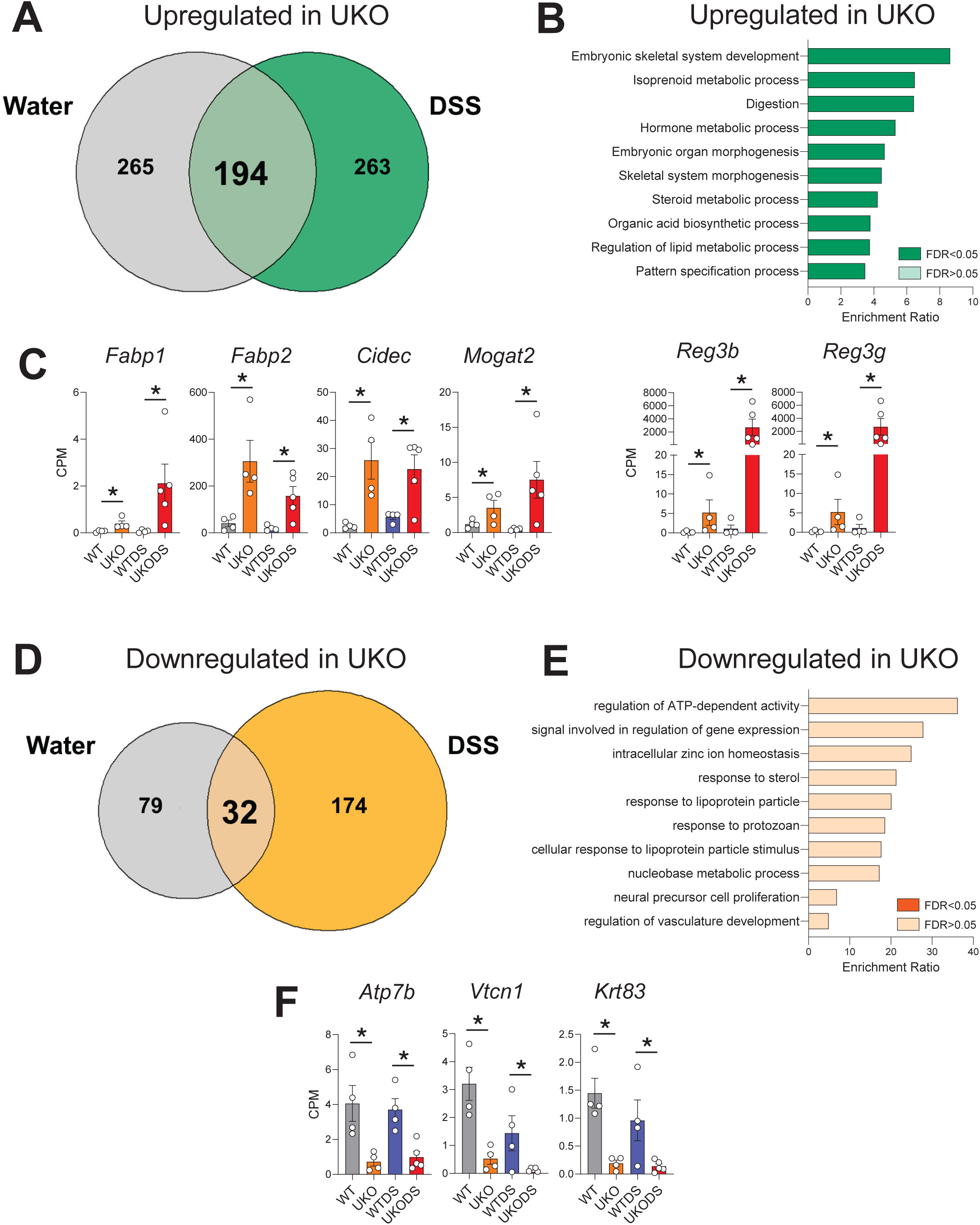
Shared transcriptional alterations in the colon of Ube4A knockout mice at baseline and during colitis. Bulk RNA-seq analysis comparing gene expression in UKO versus WT colons under both unchallenged (water) and DSS-treated conditions. **(A)** Venn diagram, **(B)** biological pathways analysis, and **(C)** single representation of genes consistently upregulated in UKO vs WT colons across both basal (WT, UKO) and DSS (WTDS, UKODS) conditions. **(D)** Venn diagram, **(E)** biological pathways analysis, and **(F)** single representation of genes consistently downregulated in UKO vs WT colons across basal and DSS conditions. Data are presented as mean ± SEM. Each dot represents one individual mouse (n=4-5/group). *P < 0.05 by Benjamini-Hochberg.

## DISCUSSION

In this study, we identify Ube4A as a previously unrecognized regulator of colonic homeostasis, revealing that its loss predisposes the colon to exaggerated inflammatory responses. Using an integrated approach combining genetically engineered models, histopathology, transcriptomics, and immunophenotyping, we demonstrate that global deletion of Ube4A markedly increases susceptibility to DSS-induced colitis in mice. Upon inflammatory challenge, UKO mice exhibited greater weight loss, higher disease activity, shortened colon length, and heightened mucosal damage compared to controls. The worsened disease severity was accompanied by a substantial increased inflammatory gene expression and a loss of mucus-producing goblet cells, well documented in human IBD [42], indicating an impaired ability to prime an appropriate immune response and maintain epithelial barrier defenses under inflammatory stress. Although unchallenged Ube4A-deficient mice appeared grossly normal, with no overt histological abnormalities, their colons displayed extensive basal transcriptional remodeling, pointing to activation of lipid metabolic and stress-adaptive programs that precede inflammation. These data suggest that Ube4A sustains a transcriptional program that supports epithelial metabolic balance and protects against a “pre-stressed” epithelial state even in the absence of overt inflammation.

During DSS-induced colitis, Ube4A-deficient mice mounted amplified inflammatory responses, characterized by heightened expression of acute inflammatory mediators, together with a pronounced type-2–skewed response. A prominent feature of this response was the accumulation of mast cells, highly expressing proteases such as MCPT-1. Consistent with mast-cell–driven tissue injury, UKO colons also showed upregulation of several matrix metalloproteinases (MMPs) including Mmp3, 10 and 13, which have been all reported to be upregulated in human IBD [43–46]. Due to their proteolytic activity, MMPs are well-established inducers of extracellular matrix degradation and epithelial erosion during inflammation and can also regulate the activation and processing of inflammatory mediators [47, 48]. Prior work has also shown that mast cells contribute to intestinal tissue injury in several models of IBD, in part through protease release and induction of MMPs [49]. Indeed, trials targeting mast cell tryptases have demonstrated therapeutic potential in both human and animal models of colitis [50, 51]. However, other studies indicate that mast cells may support epithelial repair during recovery phase through cytokine-dependent interactions with damaged epithelium [52], suggesting that mast cell activation during inflammation may exert context-dependent effects. Together, our findings raise the possibility that Ube4A influences mast-cell–associated inflammatory responses, although the underlying mechanisms and functional significance will require further investigation.

To gain insight into molecular determinants of heightened susceptibility to colitis, we examined colonic homeostasis in untreated Ube4A-deficient mice by RNA-seq. Our data show that, even at baseline, UKO colons exhibited a high enrichment of transcriptional pathways involved in lipid and isoprenoid metabolism, together with elevated expression of epithelial stress- and damage-responsive genes. These findings align with a growing body of evidence indicating that dysregulated lipid handling including uptake, storage, and export, within the intestinal epithelium is closely linked to mucosal inflammation in IBD [53, 54]. Colonic epithelial cells normally rely heavily on fatty acid oxidation (FAO), particularly of microbiota-derived short-chain fatty acids, as a primary source of energy. Shifts away from FAO toward glycolysis are increasingly recognized as a hallmark of inflamed IBD mucosa [27, 55]. Recent studies using organoid models, have shown that epithelial cells derived from patients with pediatric ulcerative colitis display hypermetabolism, mitochondrial stress, with enhanced lipid accumulation and upregulation of lipid hydrolysis and transport pathways [56]. Importantly, these metabolic changes are coupled to increased chemokine production and neutrophil chemotaxis, which can be mitigated by inhibiting PPARα [56]. Our findings are consistent with recent work demonstrating that the E3 ubiquitin ligase Hakai regulates fatty acid synthase (FASN) stability through ubiquitination-dependent degradation, thereby controlling FASN-mediated lipid accumulation, a process linked to the development of IBD [22]. Defects in lipid export, rather than lipid accumulation alone, are closely associated with disease severity in IBD. Clinical studies show that abnormalities in systemic lipid handling, particularly reduced HDL cholesterol and apolipoprotein A1, key mediators of lipid efflux, correlate with more aggressive disease [57]. Consistent with these findings, additional reports documented abnormal serum lipid profiles in patients with active or complicated IBD, supporting the concept that impaired lipid export and lipid imbalance modify disease course rather than simply reflecting chronic inflammation [58, 59]. Although these clinical studies do not pinpoint the cellular compartments in which lipid dysregulation is most consequential, our data suggest that Ube4A plays a critical role in maintaining colonic metabolic balance. Its loss may induce epithelial stress potentially lowering the threshold for heightened inflammatory responses following injury.

Human datasets provided further support for the relevance of these findings. Analysis of colonic scRNA-seq data showed that *UBE4A* gene expression is enriched in epithelial cells, particularly in proliferating transit-amplifying progenitors that give rise to goblet cells [31]. We found that *UBE4A* expression progressively declined from healthy colon to non-inflamed IBD tissue and was lowest in inflamed biopsies, mirroring the epithelial stress signatures and goblet cell depletion observed in Ube4A-deficient mice. Prior clinical studies reporting elevated anti-Ube4A autoantibodies in Crohn’s disease, especially in patients with more severe or complicated disease [24], further suggest that perturbations in Ube4A may be linked to epithelial dysfunction and immune dysregulation in human IBD. While the functional consequences of these autoantibodies are not yet known, our findings provide a mechanistic context in which altered Ube4A expression or activity could contribute to disease pathogenesis and/or progression.

Our study presents some limitations. Because this study used a global knockout model, we could not determine whether Ube4A acts primarily in epithelial, stromal, or immune compartments. Further studies using tissue-specific deletion models will be required to dissect these contributions. Additionally, our DSS model primarily reflects acute epithelial injury rather than chronic immune-mediated inflammation, and complementary models will be necessary to fully define Ube4A’s role across the course of IBD pathology. Finally, the molecular substrates and pathways through which Ube4A regulates epithelial stress and mucosal immunity remain to be elucidated.

In summary, our work defines Ube4A as a new gatekeeper of epithelial stress resilience and colon inflammation. Loss of Ube4A induces a latent epithelial stress state, associated with upregulation of lipid metabolic programs that may compromise mucosal barrier defenses during injury, and amplify mast-cell–driven inflammation during experimental colitis. Alongside human data linking reduced *UBE4A* expression and dysregulated lipid metabolism to more severe IBD phenotypes, these results support a model in which Ube4A safeguards a metabolically balanced epithelial state that protects against inflammatory triggers. Overall, this study provides a foundation for future work aimed at elucidating how Ube4A-dependent ubiquitin signaling integrates metabolic and inflammatory stress to preserve mucosal homeostasis.

## MATERIAL AND METHODS

### Mice and treatment

All animal studies were approved by the Saint Louis University Institutional Animal Care and Use Committee. Mice on C57BL6 background (JAX) were housed with a maximum of five mice per cage in a controlled atmosphere at 23 °C on a 12-hour light/dark cycle. Animals had free access to food and water, and their health status was monitored throughout the duration of treatment. Age-matched littermates were used in all experiments. Wild-type (WT) and Ube4A knockout (UKO) male (∼13-week-old) were used in all experiments. DSS salt (colitis grade; molecular weight, 40,000–50,000 daltons) was purchased from Thermo Scientific (Cat# J14489-22 MP Biomedicals, Irvine, CA). DSS salt was given to mice as a 2.5% solution in drinking water, ad libitum, for five consecutive days to induce acute colitis (injury phase). DSS-containing water was refreshed every 2-3 days. For functional, histological or molecular studies of the injury phase, mice were sacrificed for sample collection at the end of the five-day DSS treatment period. For analysis of body weight and disease activity index (DAI), after five days of DSS, mice were allowed to recover on normal drinking water for 15 consecutive days (recovery phase). The body weight measurement and stool collection of each mouse was conducted every day for the duration of the study (20 days). DAI was assessed based on the following scoring system: normal stool consistency with negative hemoccult: 0; soft stools with positive hemoccult (Beckman Coulter): 1; very soft stools with traces of blood: 2; watery stool with visible rectal bleeding: 3, as previously reported (14).

### Histological and immunohistochemical analyses

Colons were harvested in unchallenged mice or after five days of DSS treatment. A proximal region (∼1 cm) was cut, fixed in 4% paraformaldehyde, paraffin-embedded and 5 µm thin sections were prepared for histological analyses. Colon sections were stained for Hematoxylin and Eosin (H&E) or Alcian Blue by the SLU Advanced Spatial Biology and Research Histology Core Facility. Sections were imaged using a Cytation C10 Confocal Imaging Reader (BioTek). For the DSS treatment, the extent of colonic damage was assessed using H&E staining and scored by a blinded evaluator based on a modified scoring method adapted from [60, 61], as described in **Supplementary Table 1**. For quantification of goblet cell depletion, scanned images from alcian blue staining were converted to RGB color, subjected to color deconvolution and 10 randomly chosen different colon areas/sample were selected for measurements using the ImageJ software (2.16.0/1.54p).

### Immunofluorescence

Paraffin-embedded colon sections were subjected to heat-induced antigen retrieval in Diva Decloaker buffer (Biocare Medical) for 18 minutes at 95°C. The slides were incubated at RT for 1h in a blocking buffer (5% BSA, 0.5 % triton in PBS). The sections were then incubated overnight at 4°C with the primary antibody Anti-MCPT-1 (eBioscience cat #14 5503-82, clone RF6.1), washed in PBS-Tween 0.01% and incubated at RT with the secondary antibody. Slides were mounted using EverBriteTM mounting medium (Biotium, 23002) containing DAPI. Tissues were imaged using a DragonFly confocal microscope (Leica Microsystems, Germany) and processed using ImageJ software (2.16.0/1.54p).

### Bulk RNA-Sequencing and quantitative RT-PCR (qRT-PCR) studies

Total RNA was isolated from frozen unchallenged and DSS-treated colon samples by using QIAzol lysis reagent and the RNeasy mini kit (Qiagen, Valencia, CA) in combination with an RNase-free DNase Set (Qiagen). Total RNA integrity was determined using Agilent 4200 Tapestation. Library preparation was performed with 5ug of total RNA with a RIN score greater than 8.0. Ribosomal RNA was removed by poly-A selection using Oligo-dT beads (mRNA Direct kit, Life Technologies). mRNA was then fragmented in a reverse transcriptase buffer and heated to 94°C for 8 minutes. mRNA was reverse transcribed to yield cDNA using SuperScript III RT enzyme (Life Technologies, per manufacturer’s instructions) and random hexamers. A second strand reaction was performed to yield ds-cDNA. cDNA was blunt ended, had an A base added to the 3’ ends, and then had Illumina sequencing adapters ligated to the ends. Ligated fragments were then amplified for 14 cycles using primers incorporating unique dual index tags. Fragments were sequenced on an Illumina NovaSeq X Plus using paired end reads extending 150 bases targeting 30M reads per sample. Raw data fastqs were generated using Illumina’s bcl2fastq software. Reads were aligned to the Ensemble v101 primary DNA assembly genome using STAR RNA-seq aligner v2.9.9a1 [62]. Raw gene level counts were quantified using RSubread-featureCounts [63]. Raw counts matrices were normalized using the Trimmed Mean of M-Values (TMM) method. Subsequent differential gene expression testing was performed using the R edgeR package [64]. After fitting the data using a negative binomial generalized linear model, differentially expressed genes were tested for using quasi-likelihood (QF) F-tests. Genes were filtered by Benjamini-Hochberg analysis and those with a raw p-value < 0.05 and a log(fold change) below −1.5 or above 1.5 were considered to be significantly differentially expressed. Over-representation analysis (ORA) was performed on differentially expressed gene lists using Webgestalt [65]. qRT-PCR analyses were performed following a standard, SYBR green-based, ΔΔCt method, using our previously published protocol [26]. The list of primers used is provided in **Supplementary Table 2.**

### Flow cytometry

Fluorochrome-conjugated antibodies were purchased from Biolegend (anti-CD45, cat. #157612, clone QA17A26; anti-CD8a, cat. #100750, clone 53-6.7; anti-CD4, cat. #100555, clone RM4-5; anti-CD19, cat. #115546, clone 6D5; anti-CD11b, cat. #101222, clone M1/70; anti-NK1.1, cat. #108710, clone PK136; anti-CD117, cat. #135115, clone BL; anti-IL-33Ra(ST2), cat. #145309 clone DIH9), and Invitrogen (anti-FCeR1 alpha, cat. #12-5898-82, clone MAR-1). All flow cytometry verified antibodies were used at a 1:200 dilution and all intracellular flow cytometry verified antibodies were used at a 1:100 dilution. Near-IR fluorescent reactive dye (live/dead) was purchased from Invitrogen (cat. #L34975A) and used at a 1:1000 dilution, and Fc block was purchased from Biolegend (cat. #101320) and used at a 1:200 dilution. Intracellular staining of nuclear-associated proteins was performed using the eBioscience cellular permeabilization kit (cat. #00-5523-00) per the manufacturer’s instructions. Briefly, cells were processed and stained *ex vivo* with live/dead and cell surface markers described above. Cells were then fixed, permeabilized, and stained with antibodies specific for the intracellular proteins T-bet (Biolegend; cat. #644824, clone BL), Gata3 (Invitrogen; cat. #15-9966-42, clone TWAJ). Flow-cytometric analysis was performed on LSRFortessa Cell Analyzer (BD Biosciences) in the Saint Louis University Flow Cytometry Core Facility, and data analyzed using FlowJo v.10 software (Tree Star Inc.).

### Single cell RNA-seq data mining in human colon

The data from Smillie et al., [30] was downloaded from the Single Cell Portal (accession number: SCP259). Processed data were used according to the authors’ instructions provided at https://github.com/cssmillie/ulcerative_colitis?tab=readme-ov-file. Epithelial, immune and stromal single cell objects were analyzed using the *UpdateSeuratObject* function of the Seurat package (version 5.0.0) [66] in R.

### Statistical analysis

All statistical analyses were performed using Prism 7 (GraphPad Software, San Diego, CA) and all data were presented as Mean ± SD or SEM. Differences between groups were significant when P < 0.05, using two-way ANOVA, Mann-Whitney, Wilcoxon rank sum test, or unpaired Student t-test.

## ACKNOWLEDGMENTS

This study was supported by Saint Louis University Startup fund, seed grant from the SLU Department of Pharmacology and Physiology and President Research Fund to Vincenza Cifarelli, and Saint Louis University and University of Florida Startup funds and R01DK132162 to Anutosh Chakraborty. We are grateful to Jay McQuillan, Kjirstin Osland, Atika Malik and Shikara Patel for technical assistance. We also acknowledge SLU Flow Cytometry Core, and the Genomics and Bioinformatics Core for assistance with experiments.

## AUTHOR CONTRIBUTIONS

AC and VC conceived, designed and supervised the studies. SG, MC, SGN, AV, EE, RMT, AC and VC conducted the studies. SG, SGN, DDB, MB, EE, RMT, AC and VC performed sample analyses. SG, MC, SGN, AV, AC and VC analyzed the data. SG, SGN, DDB, MB, AC and VC performed the statistical analyses. SG, AC and VC wrote the manuscript. AC and VC obtained funding for the work. AC and VC are the guarantor of this work and, as such, had full access to all the data in the study and takes responsibility for the integrity of the data and the accuracy of the data analysis. All authors critically reviewed and edited the manuscript.

---

**Supplementary Figure 1.**
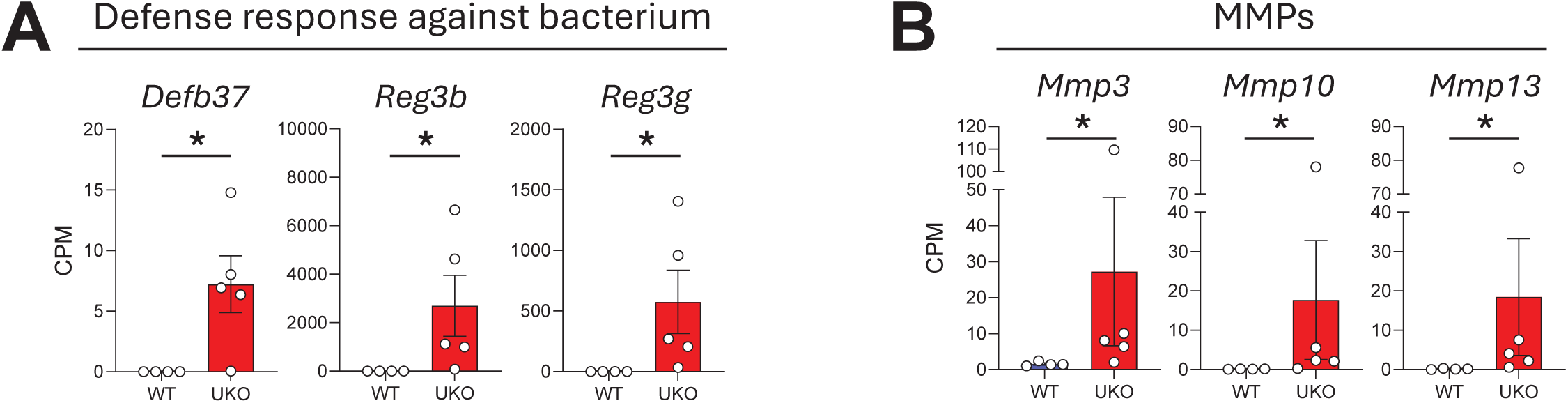
Relative colonic expression of genes related to **(A)** defense response against bacterium and **(B)** matrix metalloproteinases (MMPs), analyzed by bulk RNA-seq in five-day DSS-treated UKO and WT mice. Data are presented as mean ± SEM. Each dot represents one individual mouse (n=4-5/group). *P < 0.05 by Benjamini-Hochberg.

**Supplementary Figure 2.**
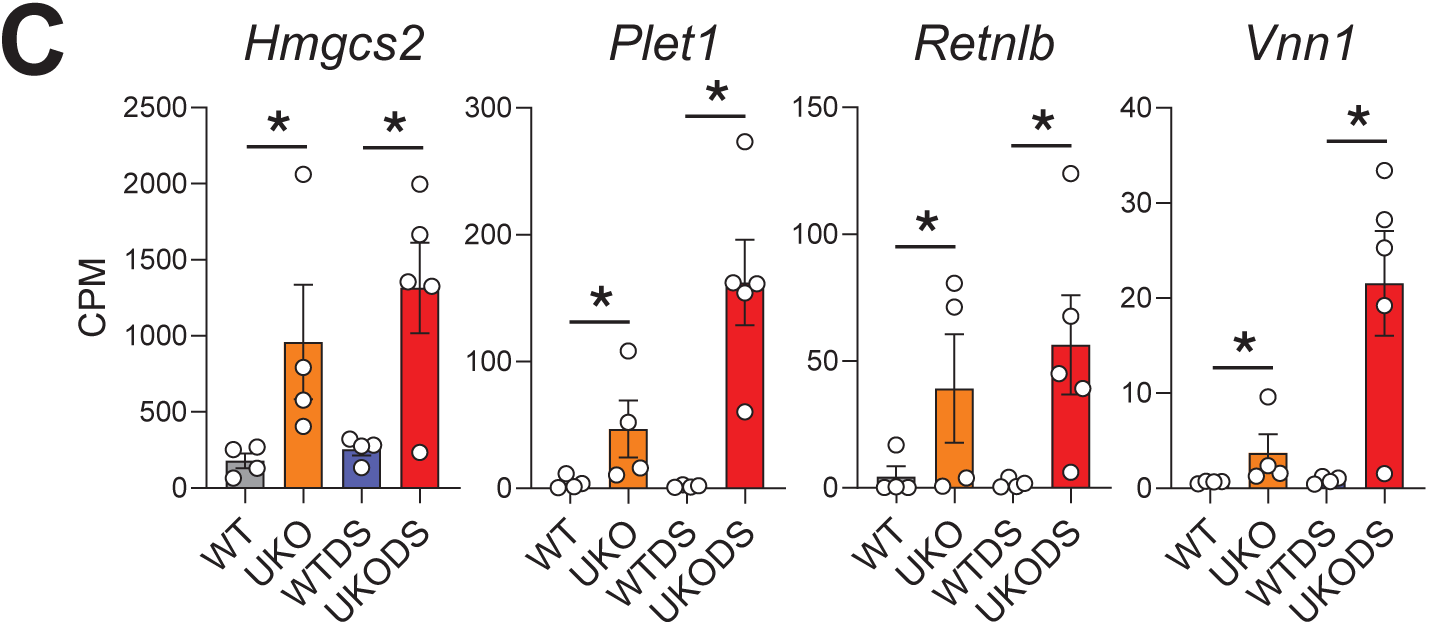
Relative colonic expression of genes related to epithelial stress or damage between WT and UKO mice across both basal (WT, UKO) and DSS (WTDS, UKODS) conditions. Data are presented as mean ± SEM. Each dot represents one individual mouse (n=4-5/group). *P < 0.05 by Benjamini-Hochberg.

**Supplementary Table 1.** Experimental colitis scoring criteria.

| Category | Feature | Score value | Criterion |
| --- | --- | --- | --- |
| Degree of Immune cell infiltration |  | 0 | absence |
|  |  | 1 | mild |
|  |  | 2 | moderate |
|  |  | 3 | severe and transmural infiltrate |
| Edema in the submucosa |  | 0 | absence |
|  |  | 1 | mild |
|  |  | 2 | moderate |
|  |  | 3 | severe |
| Extent of muscle thickening |  | 0 | absence |
|  |  | 1 | moderate thickening (mostly external muscularis) |
|  |  | 2 | extensive muscle thickening (both external and mucosae muscularis) |
| epithelial architecture and damage | Crypt architecture | 0 | normal epithelial architecture |
|  |  | 1 | mild decreased crypt density or presence of crypt hyperplasia |
|  |  | 2 | moderate crypt distortion with <50% decreased crypt density and/or severe hyperplasia |
|  |  | 3 | severe crypt distortion with >50% loss of entire crypts |
|  | Goblet cell loss | 0 | absence |
|  |  | 1 | mild: <20% |
|  |  | 2 | moderate: 30-50% |
|  |  | 3 | severe: >50% |
|  | Cryptitis (immune infiltrate between the crypts) | 0 | absence |
|  |  | 1 | presence |
|  | Erosion (loss of surface epithelium) | 0 | absence |
|  |  | 1 | focal site(s) of erosion |
|  |  | 2 | diffuse and extensive erosion |
|  | Ulceration (mucosal injury reaching beyond muscularis mucosae) | 0 | Absence |
|  |  | 1 | Presence |

**Supplementary Table 2.**
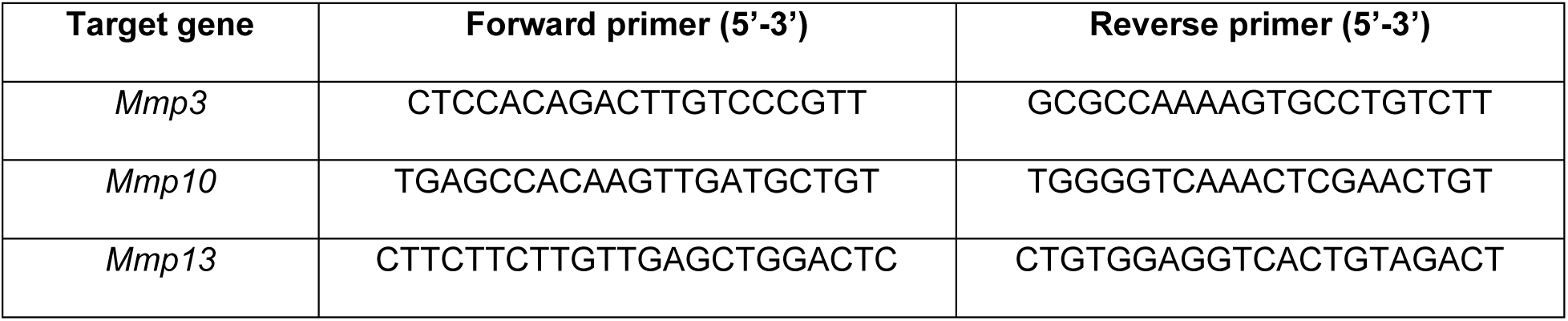
List of primers for qRT-PCR analysis.

